# Topology in Motion: Geometry-Driven Defect Dynamics in Social Wasp Nests

**DOI:** 10.1101/2025.08.31.673251

**Authors:** Tarini Hari, Somendra M. Bhattacharjee, Shivani Krishna

**Affiliations:** Department of Biology, Trivedi School of Biosciences, Ashoka University, Sonepat 131029, India; Department of Physics, Ashoka University, Sonepat 131029, India

## Abstract

Emergence of order in social insects is exemplified by their nesting architectures. Paper wasps construct hexagonal nests that, like crystals, harbour topological defects. In a paper wasp nest, we identify transient defects that migrate by local wall reorientation and vertex addition, akin to dislocation glide. Across growth phases, non-hexagonal polygons undergo short-range transformations in form and position. The transitions resemble alternating *Y–*Δ *transformation–node release* and *release–and–straighten events*, familiar from electrical networks. Burgers circuits show that passage through intermediates does not reduce the distortion in the nest, and all transitions conserve topological charge, thereby explaining the coupling of non-hexagonal cells. These dynamics propagate defect motion only to a finite extent, after which the defect stabilizes without global reorganization or functional gain. Our findings demonstrate that social insect nest building naturally realizes dynamic topological processes, shaped by material properties, construction rules, and geometric constraints.

---

Biological assemblies, ranging from epithelial tissues, viral capsids, to bird flocks, fish schools, and insect nests, display remarkable order, often emerging from simple, local interactions among many agents ^1–4^. Social insect nests exemplify this, with paper wasps building hexagonal cells from plant fibres that optimize packing and strength ^5–7^, but like crystals, harbour topological defects such as pentagons and heptagons^8,9^. Honeybees can reshape wax cells to correct or exploit such non-hexagonal cells^10–12^, while the hardened plant fibre walls of wasp nests constrain repair, making it more challenging to deal with them. These topological defects in honeycomb lattices can take the form of disclinations, such as a sole pentagon or heptagon, dislocations such as pentagon–heptagon (7–5) pairs, higher order defects like Stone-Wales defects ^9^. A single pentagon or heptagon embedded in a hexagonal lattice, such as the nest, disrupts the local orientational order and introduces intrinsic curvature, typically requiring the lattice to deform into the third dimension. However, when a 5-sided and a 7-sided cell are placed adjacent to each other, their opposite angular distortions cancel, preserving the overall planarity of the nest structure. This 7-5 pair forms a defect dipole (known as a dislocation), which differs from a disclination, that affects the translational order of the nest.

While topological defects and their dynamics have been extensively studied in a wide range of ordered systems, including cosmological contexts ^13^, their emergence and dynamics in biological architectures remain largely unexplored. Here, we report an emergent topological defect state, namely a 7–4–7 triplet, in a paper wasp nest. This defect appears as an intermediate configuration that facilitates the shift of a dislocation within the nest through purely geometric rearrangements.

The geometry-driven dynamics of such a defect state have not been previously documented in insect-built architectures.

The study was conducted on *Polistes wattii* species during the onset of their summer nesting. *P. wattii* is a primitively eusocial species with a widespread distribution (Asia and Middle East). They are known to have two nesting cycles, one in the spring (March to May) and another in the summer (June to October) ^14^. Wasp nests were imaged 2–3 times daily from a distance of 15 m. Tracking the nest growth trajectory of one of these nests revealed the construction of a topological defect. This nest expanded from 40 to 258 cells during the period of observation. Nest expansion was divided into three phases to illustrate the progression of the non-hexagonal complex. To analyze local rearrangements, the nest was treated as a planar network, with walls as edges and vertices as nodes. The sequence of construction and rearrangement is detailed below in three distinct phases: initiation, migration, and stabilization.

## Phase I-Initiation

In the initial phase, when the nest comprised of 35 cells, we noted a pentagon–heptagon pair at the nest margin (Fig. 1.1). This pentagon was transformed into a hexagon, facilitated by the emergence of a quadrilateral (via the addition and removal of vertices). Integration of this quadrilateral to the nest through its peripheral growth resulted in a second heptagon, thus forming a 7-4-7 cell complex (Fig. 1.2). The emergence of this quadrilateral is an interesting phenomenon. The vertex supporting three edges has a Y-shaped configuration, and we predict that a Y-Δ transformation converts this into a triangular loop of three edges with three vertices in a Δ-shape. With a pentagon as a defect, such a transformation enables the addition of a vertex to transform the pentagon into a hexagon (marked as 1, Fig. 1.3). Concurrently, an adjacent vertex undergoes a Y-Δ transformation to create two converging triangles with a shared vertex. Since only vertices with three-points are preferred by wasps, the four-point vertex is eliminated to form two edges whose straightening leads to the formation of a quadrilateral. The process locally accumulates material, observed as thickened walls in some of the cells after completion. The Y-Δ mechanism was first described in electrical network theory (the star–triangle transformation) and has been widely used for understanding planar lattice models in statistical mechanics^15^.

**Figure 1:**
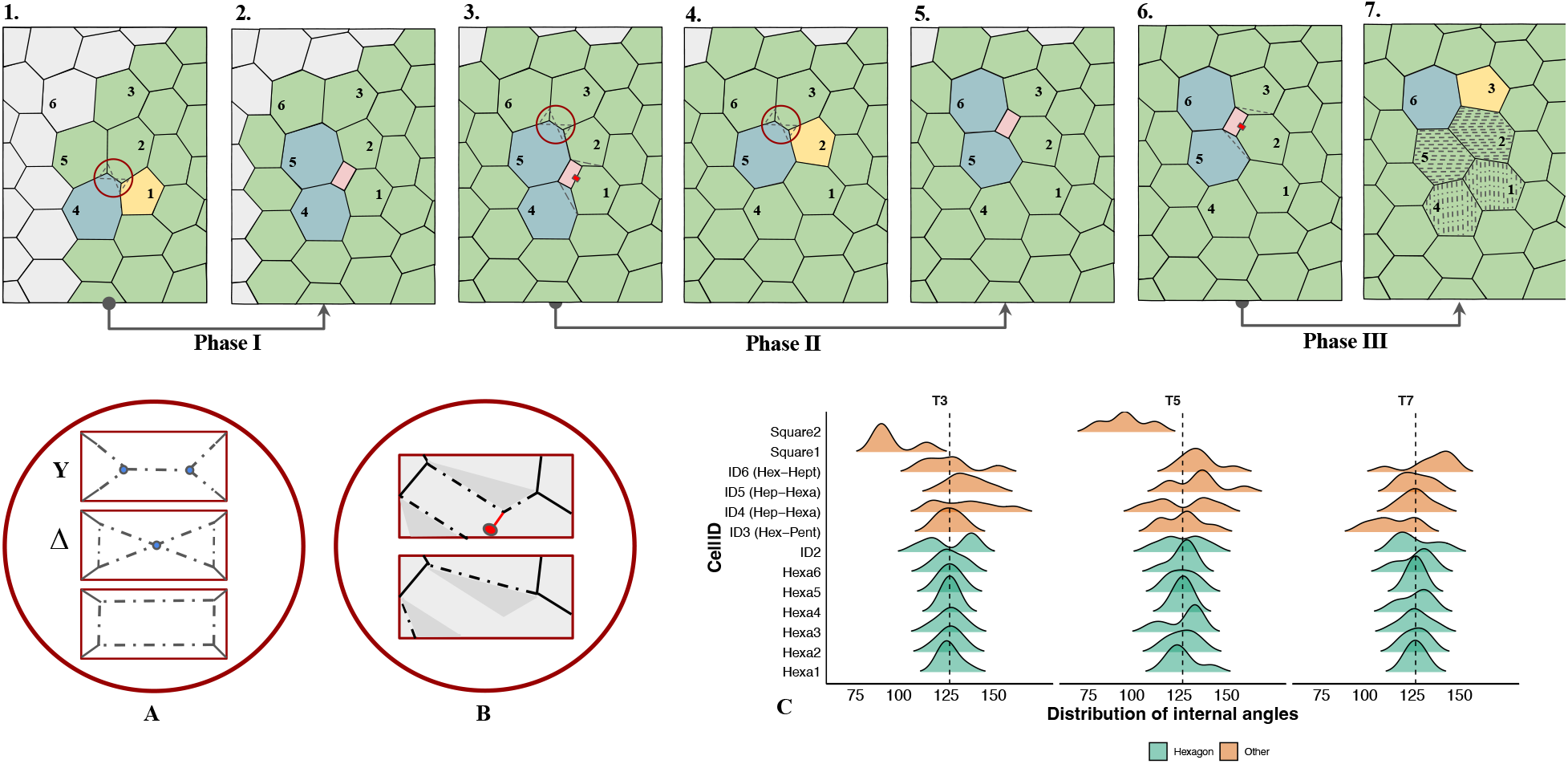
A schematic representation of topological defects, their positions and changes over time. **(1)** Phase I: Topological defect region being constructed (pentagon-heptagon pair at the nest margin) at a 40-celled stage. **(2)** Phase I: Emergence of the 7-4-7 dislocation. **(3**,**4**,**5)** Phase II: Migration of 7-4-7 complex across the nest. Box 4 illustrates the possibility of transformation to a 7-5 pair in Phase II. **(6**,**7)** Phase III: Transformation of the 7-4-7 to a stable 7-5 defect state. Shaded cells mark cells that were previously non-hexagons, indicating the path of the defect motion. Panels **(A** and **B)** Expanded view of the circular regions indicating the Y-Δ transformations-node release, release-and-straighten events resulting in the quadrilateral. **(C)** Ridge line plot: The density distribution of internal cell angles of hexagonal (green, Hexa1-6 and ID2) vs non-hexagonal (orange) are plotted across time points (T3, T5 and T7 represent Phases I, II and III, respectively) of migration. Quadrilaterals are shown as Square1 and Square2; ID4 and ID5 were cells that were heptagons transformed to hexagons; ID6 was a hexagon that was converted to a heptagon and ID3 was a hexagon that was converted to a pentagon.

## Phase II-Migration

During this phase, the nest grows from 55 cells to a 125-cell stage. Phase II marks the stage where the spatial migration of the 7-4-7 defect within the nest has been identified. Changes in the internal wall angles of the non-hexagonal cell complex indicate a transient state, serving as a precursor to a translational shift. The distribution of interior angles for cells that remained hexagonal (mean *±* sd = 120 *±* 6.3 degrees) differed from cells that experienced transformations (e.g., ID4, ID5 in Fig. 1C; range of mean angles = 90-129 degrees). Migration was found to occur incrementally, one cell at a time, with the annihilation and subsequent reconstruction of another quadrilateral hinging on a pivot cell (marked as 5 in Fig. 1C). This pivot cell retained its defect state through the process. The annihilation of the quadrilateral on one face of the pivot cell occurred via a release-and-straighten, which involves an edge cut opposite to the pivot cell (‘release’), followed by straightening of the adjacent edges (‘geometric relaxation’). Its subsequent construction on a different face was possibly a repetition of the Y-Δ transformations-node release events as during Phase I. The removal or addition of the quadrilateral also transformed a neighboring heptagon into a hexagon (cell-4), and vice versa (cell-6), completing the one-unit displacement of the entire 7-4-7 defect. If the quadrilateral was not rebuilt, a pentagon-heptagon (7–5) pair could have emerged at this stage (indicated in Fig. 1.4), signaling a change in defect type rather than its position. The observed construction of a second quadrilateral hinted towards a choice made at this stage to continue the translational mobility upwards (in the direction of the nest growth) as an attempt towards ‘ironing’ out non-hexagonal cells.

## Phase III-Stabilization

In Phase-III, a similar sequence of transformations as seen in Phase II was observed without the last step of reconstruction. Readjustments to internal wall angles stretch the quadrilateral before being annihilated. Removal of a vertex from the adjacent cell (marked as 3, Fig. 1.7) converts it to a pentagon, while the pivot cell (marked as 6, Fig. 1.7) retains its heptagonal form. A resultant 7-5 pair is formed two cells away from the Phase-I pentagon location. This state remains consistent for the rest of the nest’s growth trajectory. Fig. S1 provides a detailed sequential overview of nest growth, depicting the progressive changes in the positions of the defect complex across growth phases.

The persistence of the 7–5 cell pair, despite repeated local angle readjustments, indicates that defect motion reaches a locally accommodated endpoint without global reorganization, motivating the arguments below on the limits and possibilities of emergence of defect complexes in such nests. Local changes alone cannot repair a topological defect without inducing global change to a system, the latter not being an easy possibility in dried paper nests. Additionally, given the relatively small size of the quadrilateral, no functional advantage for brood care or storage is evident (in honeybee nests quadrilaterals were used as filler cells^10^), and therefore we predict that their appearances are incidental and not adaptive. The three phases demonstrate an apparent local motion of the topological defect, exhibiting transformations in form and spatial position only in the short range.

This informs our proposition that the alternating *Y-*Δ *transformations-node release* and *release-and-straighten* events can propagate the defect’s movement, at the expense of removal of walls and the construction of new walls.

To evaluate whether the magnitude of distortion provides cues for this displacement, we computed the Burgers vector, which represents the *net lattice translation* around a closed circuit enclosing the dislocation (the circuit was drawn manually by tracing an equal number of cell centers in each direction; in a defect-free region, the loop closes with zero excess). The Burgers vector associated with the dislocation was comparable in both Phase II (7-4-7) and Phase III (7-5) configurations (Fig. 2). Thus, the construction of a relatively complex defect state does not appear to confer the advantage of reduced distortion. This invariance arises from the topological character of the defects, specifically the relationship between dislocations and pairs of disclinations (known as a disclination-dipole). The dipole direction, defined as the vector from the center of the 7-sided cell to the center of the 5-sided cell, determines the orientation of the dislocation. The associated Burgers vector lies in-plane and is perpendicular to this dipole. In more complex configurations, such as the 7-4-7 defect, the structure contains two dipoles, each formed by a 7-4 pair. The vector sum of these dipoles (of zero total charge) results in a net dipole aligned with that of the original 7-5 pair. Consequently, the transformation 7-5*↔*7-4-7 preserves the Burgers vector, and maintains the global topological character of the nest (Fig. 2).

**Figure 2:**
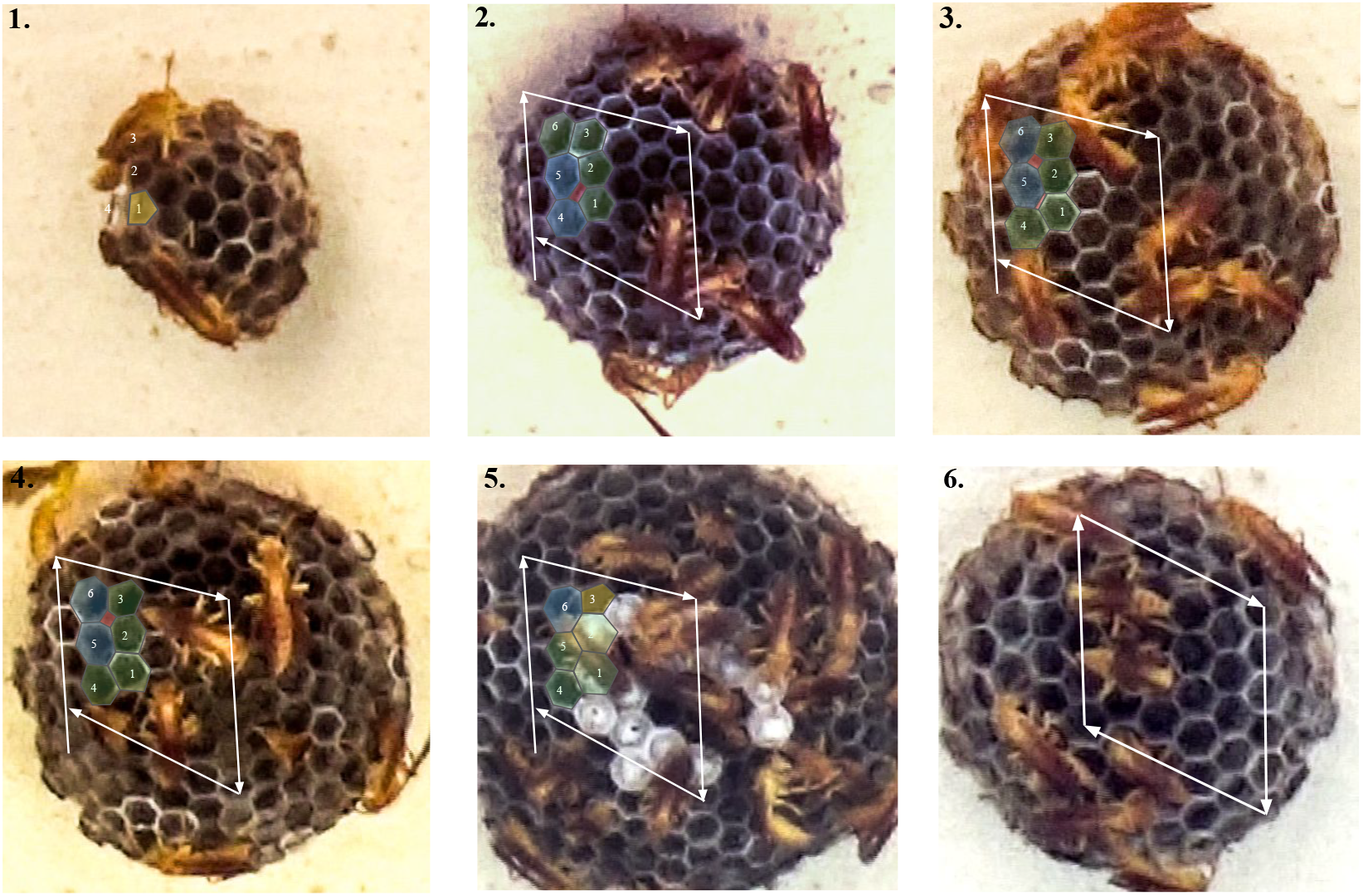
Burgers circuit at various stages of nest growth depicting the vector with additional displacement in the defect region, in contrast to the region without defect where the loop closes without an excess. **(1)** Initial pentagon visible during Phase I. **(2)** The Burgers vector drawn around the 7-4-7 complex before migration, indicating the extra step. **(3)** The vector around the 7-4-7 region during migration via annihilation of a quadrilateral and emergence of another. **(4)** The Burgers vector for the 7-4-7 complex post-migration, one unit cell away from the beginning of Phase II. **(5)** The Burgers vector for the stabilized 7-5 defect during Phase III, located two unit cells away from initialization in Phase I. **(6)** The Burgers vector of the hexagonal lattice without defects (enclosing hexagonal cells).

Interestingly, the transformations done by wasps also reflect topological charge conservation. A hexagonal lattice (wasp nest) with six-fold rotational symmetry allows for five types of topological defects (disclinations): 3-, 4-, 5-, 7-, and 8-sided polygons, all of which have 3-point vertices. A defect carries a topological charge *q* such that a cell with fewer than six sides (such as 5, 4, or 3) carries a positive charge, while a cell with more than six sides (7, 8, 9, etc.) carries a negative charge (an overview of charges is given in Table 1). A closed loop around a disclination does not produce any Burgers vector; instead, it produces a change in the orientation of the starting hexagon by an angle given by 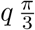. For example, +1 = (+2) + (−1), a pentagon (+1) is topologically equivalent to a quadrilateral (+2) combined with a heptagon (−1); this implies that 7-5 has the same net topological charge as a 7-4-7. Furthermore, the purported Y-Δ transformations and the resulting quadrilateral may help transition from a pentagon to an ideal hexagon. However, while respecting local topological charge conservation, these attempts to ‘iron out’ defects merely relocate charges as zero net charge cannot be achieved by local moves.

**Table 1:** Topological charge *q* = (6 *− n*) mod 6 assigned to polygonal cells in a 3-coordinated hexagonal lattice, where *n* is the number of sides. Topological charge is also known as winding number ^16,17^. It reflects the degree of distortion from ideal hexagonal order and influences the geometric orientations of the surrounding hexagonal cells.

| Polygon type | $n$ | Topological charge | Alternative |
| --- | --- | --- | --- |
| Quadrilateral | 4 | $(6 - 4) \bmod 6 = 2$ | +2 |
| Pentagon | 5 | $(6 - 5) \bmod 6 = 1$ | +1 |
| Hexagon | 6 | $(6 - 6) \bmod 6 = 0$ | 0 (no defect) |
| Heptagon | 7 | $(6 - 7) \bmod 6 = 5$ | −1 |

Topological defects with non-hexagonal cells and their occurrence as triplets and pairs have been observed in insect wings, *Xenopus* tail epithelia, and honeybee nests ^10,18^. In wasp nests, their continual wall-angle adjustments suggest that some of these defects are transient features that do not contribute to optimal architectural design. The process of defect annihilation followed by construction during Phase II resembles behaviors seen in complex liquid crystals ^19^. Properties of construction material and geometric constraints appear to drive such dynamic states in wasp nests.

These dynamics suggest a behavioral signature: if defects are being ‘ironed out’ via local Y–Δ transformations, activity of most individuals should concentrate at the defect and briefly perturb growth symmetry; to test this, we mapped wasp occupancy across the nest and calculated diameter ratios. The spatiotemporal mapping of wasps revealed nonrandom nest behaviors (quantified as time spent in each region, Fig. S2). In Phase I, new cells accumulated along defect-free margins, producing an elliptical outline. The diameter ratios deviated from 1 (range of DR = 0.87 to 1.16), indicating that defect construction transiently disrupted nest symmetry. For a perfect hexagonal nest that exhibits isotropic growth, the diameter ratio (major/minor axis or aspect ratio) is expected to have a value of 1 throughout its growth. During Phases II–III, a wasp frequently occupied the defect site, consistent with pulp deposition, inspections, and vertex-wall readjustments. Time spent at the defect region was slightly higher than the non-defect regions (mean: 66.6 s vs 41.6 s, Fig. S2). These patterns suggest regular inspection and assessment of construction at defect sites.

What drives such geometric rearrangements, and what are their consequences? Nest construction has been explained largely through template-based and stigmergy-based mechanisms^6,20^. In our study, we find indications of an interplay of both mechanisms. Our analysis showed that a non-hexagonal cell near the pedicel was ultimately transformed into a hexagon (hinting towards template-based). We hypothesize that in smaller nests (150 cells), eggs are preferentially laid in centrally located cells, and such rearrangements may serve to ensure that these core cells are hexagonal (overview of nest growth shown in Fig. S3). This may be shaped by functional constraints: pentagons and heptagons cannot accommodate eggs as snugly as hexagons, and given the limited plasticity in egg size ^21^, non-hexagonal cells are likely avoided for oviposition. Thus, geometric corrections could be a colony-level strategy to maximize the proportion of usable brood cells. In *Polistes*, where castes are not discrete, a linear dominance hierarchy is thought to structure labor and space: dominants (typically the queen) occupy the center, while workers and subordinates cluster peripherally with limited overlap ^22,23^, providing behavioral context for who undertakes such corrections.

Put simply, our findings reveal that the emergent coordination of many individuals enables the topological reorganization of ordered biological structure. How such transformations and spatial shifts are coordinated remains an intriguing avenue for future work.

## Data availability

Data used in the current study will be available from the corresponding author on request.

## Author contributions

S.K. conceived and designed the study. T.H. collected the data. T.H., S.K. and S.M.B. performed and interpreted the analysis and drafted the manuscript. All authors gave final approval for publication.

