## Supplementary Information for "Topology in Motion: Geometry-Driven Defect Dynamics in Social Wasp Nests"

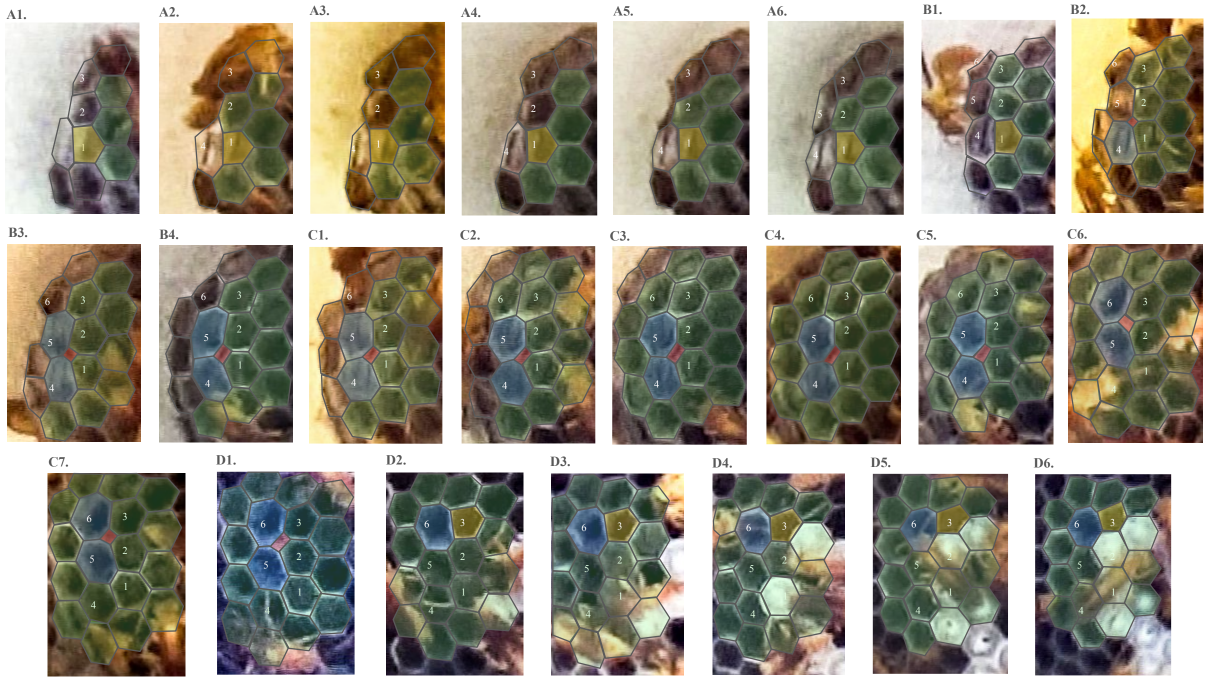


Figure S1: **(A1-6)** Topological defect region being initialized (pentagon and heptagon at the periphery of a 35-celled stage) in early Phase I. **(B1-4)** Emergence of quadrilateral as defect enters lattice, forming a 7-4-7 complex in Phase I. **(C1-4)** Visible changes in internal angles of cells along with growth in the nest in Phase II. **(C5-7)** Migration of 7-4-7 complex in Phase II. **(D1-2)** Transformation of 7-4-7 to 7-5 defect after migration, in Phase III. **(D3-6)** Stabilized 7-5 pair in Phase III. Eggs (brood caps) appear as white-filled cells in sub-panels D3-6. Hexagons are outlined in green; non-hexagons are outlined as follows-quadrilateral (red), heptagons (blue), pentagons (yellow)- overlaid on the corresponding nest images.

*correspponding author:

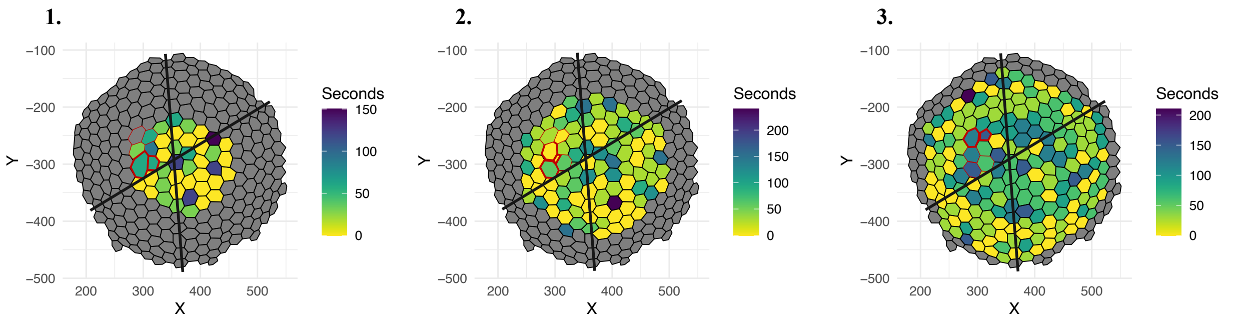


Figure S2: Time spent by wasps on the nest depicted as heatmaps. Gray cells indicate the overall size of the nest towards the end of its growth period. Darker filled colors in the cells indicates longer time spent by wasps in a cell. From recordings spanning the growth phases, we analysed snapshots of 30-s intervals in which a wasp occupied each cell. Over the growth phases, the number of wasps observed within the frame of recording varied between 4-26. For each phase, 8–10 min of sampled snapshots were aggregated to estimate occupancy. **(1)** Phase I: occupancy pattern with the initial 7–5 defect (red outline) in the upper-left quadrant. **(2)** Phase II (pre-migration): occupancy around the 7–4–7 complex (red outline) as it shifts further within the upper-left quadrant. **(3)** Phase III (post-stabilization): final occupancy pattern with the stabilized 7–5 defect (red outline farther into the upper-left quadrant).

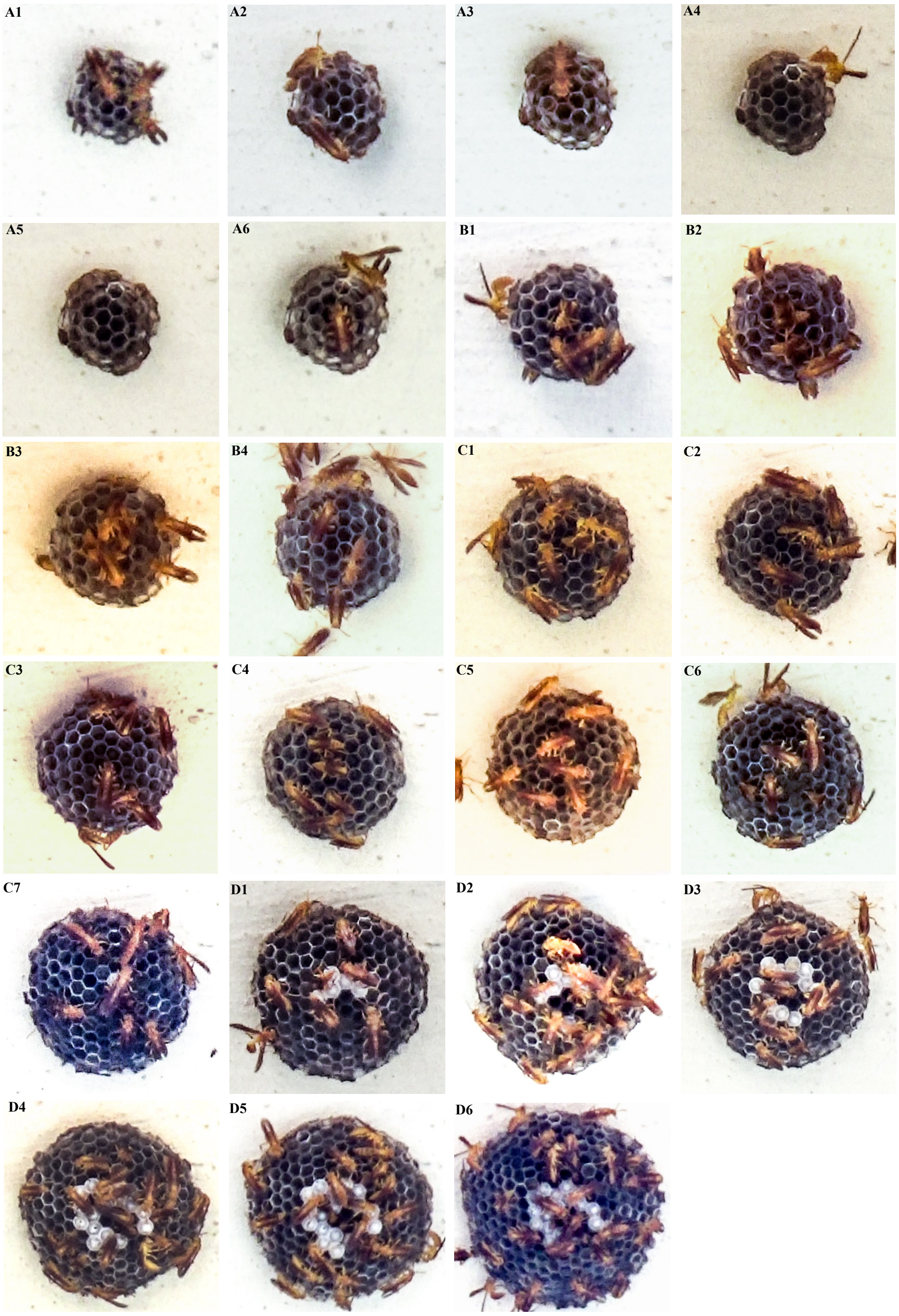


Figure S3: Sequential images documenting progressive growth of the paper wasp nest. Eggs (brood caps) appear as white-filled cells.
